# Infinitely Long Branches and an Informal Test of Common Ancestry

**DOI:** 10.1101/023903

**Authors:** Leonardo de Oliveira Martins, David Posada

## Abstract

The evidence for universal common ancestry (UCA) is vast and persuasive, and a phylogenetic test was proposed for quantifying its odds against independently originated sequences based on the comparison between one and several trees [1]. This test was successfully applied to a well-supported homologous sequence alignment, being however criticized once simulations showed that even alignments without any phylogenetic structure could mislead its conclusions [2]. Despite claims to the contrary [3], we believe that the counterexample successfully showed a drawback of the test, of relying on good alignments.

Here we present a simplified version of this counterexample, which can be interpreted as a tree with arbitrarily long branches, and where the test again fails. We also present another simulation showing circumstances whereby any sufficiently similar alignment will favor UCA irrespective of the true independent origins for the sequences. We therefore conclude that the test should not be trusted unless convergence has already been ruled out a priori. Finally, we present a class of frequentist tests that perform better than the purportedly formal UCA test.

---

Douglas Theobald [1] proposed a quantitative test to distinguish common ancestry (CA) from independent origins (IO) of a set of sequences, by modelling CA as a single tree connecting all data against two or more trees representing the IO episodes. To proceed with the actual calculations, nonetheless, a single alignment had to represent both hypotheses – which didn’t matter for the specific, highly curated data set he analysed. However, we and others have raised concerns that such a test would mistakenly infer homology (common ancestry) whenever the sequences are sufficiently similar [2, 4–7], rendering it suspicious for alignments of arbitrary quality. In particular Koonin and Wolf [2] (K&W) presented a counterexample where columns from the alignment didn’t follow any phylogenetic structure and were simply sampled from a pool of amino acid frequencies. This simulation model, called ”profile” model in [3], was enough to skew the original test into preferring UCA over the correct IO hypothesis. Theobald defended his test replying that his method would work as advertised once extended to include the true generating model of the simulated counterexamples, and also concluded that the criticisms didn’t apply for his ”very high confidence alignment” [3].

We have already shown that the test fails even for sequences simulated exactly under the described models of CA and IO, once we include the obligatory alignment optimization step [4]. We also commented on the arbitrarity of resorting to sequence similarity justifications, since all examples where his test favored IO had very low pairwise similarity themselves [4], not to mention that such a requirement would imply in a unacceptable selection bias [7].

Here we show how the K&W model was a legitimate simulation of IO, and that the UCA test fails even for a simplified version of this model where the true substitution model is amongst the tested ones. We also try to create IO alignments that satisfy the elusive constraints of quality/similarity imposed in [3] and conclude that UCA will be favored whenever the sequences are not clearly unrelated. Finally, we discuss about the lack of mathematical justification for comparing likelihoods between different alignments, and illustrate it with a simulation showing that the UCA test would fail even if we compare sequences aligned independently.

## 1 Koonin and Wolf’s profile model

K&W simulated alignments where the amino acid states for each column came from a distribution of equilibrium frequencies – that is, the state for each taxa at the *i*-th site was sampled from a discrete distribution 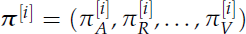. The original UCA test failed, since the log-likelihood of the whole simulated data set was always superior than the sum of the log-likelihoods of arbitrarily splitted sequences. Theobald [3] correctly pointed out that K&W’s sequences might “have evolved according to a star tree with equal branch lengths” under a MAX-Poisson evolutionary model [8], but mistakenly assumed that this was equivalent to a common ancestry scenario. The star tree from K&W model has all branch lengths equal to infinity, as we will see, which means IO. There is a key distinction between finite and arbitrarily large branch lengths, which is what ultimately discriminates UCA and IO under the original modelling (see Appendix, or [4, Supplementary Text]).

We can verify that K&W simulation corresponds to an IO scenario by using, for instance, Equation 1 of [9], which describes a similar model:

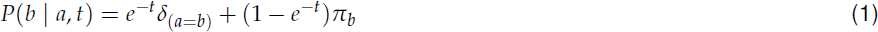

where *a* and *b* are respectively the initial and final amino acid states along a phylogenetic branch of length *t*, *δ*_*χ*_ is the indicator function^1^, and *π*_*x*_ is the equilibrium frequency of state *x*_*∈*_ (*A*, *R*, …, *V*).

As we can see, the probability of observing state *b* is influenced by the initial state *a* until *e*^-t^ *→* 0, which happens at *t* = ∞ and therefore IO. For any finite branch length *t* < ∞ the terminal states will still be correlated to the state at the root, shared among them. It’s easy to imagine that for very short branches, the state at the tips of the star tree should be very similar, since they will mostly be the same as the state at the internal node.

The source of the confusion might be that although the instantaneous substitution rate does not depend on the current state of the Markov chain, the probability of change over an arbitrary time interval does [10]. Equations (1) and (2) from [11] for example show that even for only two sequences the probability of observing state *a* in both sequences at a particular position is given by 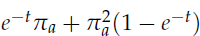 while the probability of observing state *a* in one sequence and state *b* in the other at the same column equals *π*_*a*_*π*_*b*_ (1 *− e*^*−t*^).

Therefore for small time intervals we should expect all sequences simulated under this star tree to be very similar (reflecting the common ancestry with the sequence at the root), while for longer branches they should diverge from one another until the equilibrium frequencies are reached. Under K&W’s model these probabilities are respectively 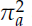 and *π*_*a*_*π*_*b*_, which are equivalent to the previous star tree values only when *e*^*−t*^ = 0, as we saw before. Therefore K&W’s model corresponds to a star tree model where all branch lengths are infinitely large – that is, the sequences are unrelated to their common ancestor.

A different question is if we can reliably estimate all parameters from the K&W simulations. The overall poor fit of MAX-Poisson as described in [8] lead us to conclude that we can’t, due to the over-parameterization being specially misleading when the number of sequences is small. There is simply not enough data to reconstruct the true frequencies vector. It might be the case that a particular data set can by chance have a corresponding phylogenetic model with finite branch lengths that explains the data equally well. But this is not the same as claiming that the data set came from such a common ancestry model.

In a nutshell, K&W’s simulations are equivalent to a MAX-Poisson model over a star tree, but with infinite branch lengths since each sequence is independent from the others. It is worth noticing that this star tree is equivalent to any other tree, or to no tree at all, due to the vanishing branches. And therefore K&W’s simulated sequences are truly originated independently, *contra* Theobald [3]. To claim otherwise would defeat, by the way, the whole phylogenetic model selection framework developed in [1]: if, for each alignment column, sampling the state (of 2 sequences or more) from a common distribution renders the data related by common ancestry, then the idea that two independent trees can represent IO would be wrong since their root positions might be two such ancestral sequences, whose columns came from an “ancestral soup” of amino acids – as we show in the Appendix.

To see it from another perspective, we can imagine any two sequences simulated by K&W as the roots of independent phylogenies. If sampling from a common pool of amino acid frequencies was enough signal for common ancestry, then the IO model (of at least one infinite branch length) would be wrong, since it does not impose restrictions on the IO evolutionary models at the root. The point is that for any combination of phylogenetic models, the IO assumption as devised in [1] is mathematically equivalent to an infinite branch connecting the nodes (apical or not). A more recent CA test explores explicitly this relation between the ancestral root states of two trees [12].

### 1.1 Our simplified simulation: a homogeneous Poisson+F

To minimize the confusion with the overly parameterized MAX-Poisson model, we reproduced K&W’s simulations but this time using a homogeneous Poisson model – that is, all columns *i* share the same equilibrium frequencies *π*^[*i*]^ = *π* = (*π*_*A*_, *π*_*R*_, …, *π*_*V*_). We were careful to include the true generating model among those tested by the model selection procedure, to be charitable and avoid misspecification issues^2^. We simulated 8 sequences with 1000 sites under a randomly sampled vector of shared amino acid frequencies. More specifically, we used INDELible [13] to simulate 2 quartets with a collapsed internal branch of length zero and all terminal branches with a huge length of 2500 – computationally equivalent to 8 independently originated sequences from a common pool of amino acids.

The AIC analysis was done with ProtTest3 (version from 18/Oct/2010) under a subset of available models, where we included the Poisson model. To be consistent with K&W we did not optimize the alignment for this analysis, although we know that the test would fail if we did align them [4]. But while K&W used a set of empirically observed amino acid frequencies to sample from, we simulated these frequencies *π* from uniform distributions – each simulated data set had a distinct frequency set, but all sites within a simulation shared the same values. We use Δ*AIC* = *AIC*(*IO*) *- AIC*(*UCA*), such that positive values of Δ*AIC* favor UCA, and a difference in AIC larger than 10 indicates that the model with larger AIC has practically no support when compared to the smaller one [14, 15]. Figure 1 shows that in most (**>** 97%) simulations the UCA was wrongly favored according to Theobald’s test, despite the large tree lengths making us suspicious about these data. Not only that, 75% of the replicates showed **very strong** support for the wrong hypothesis (that is, considering only those with Δ*AIC* **>** 10).

**Figure 1:**
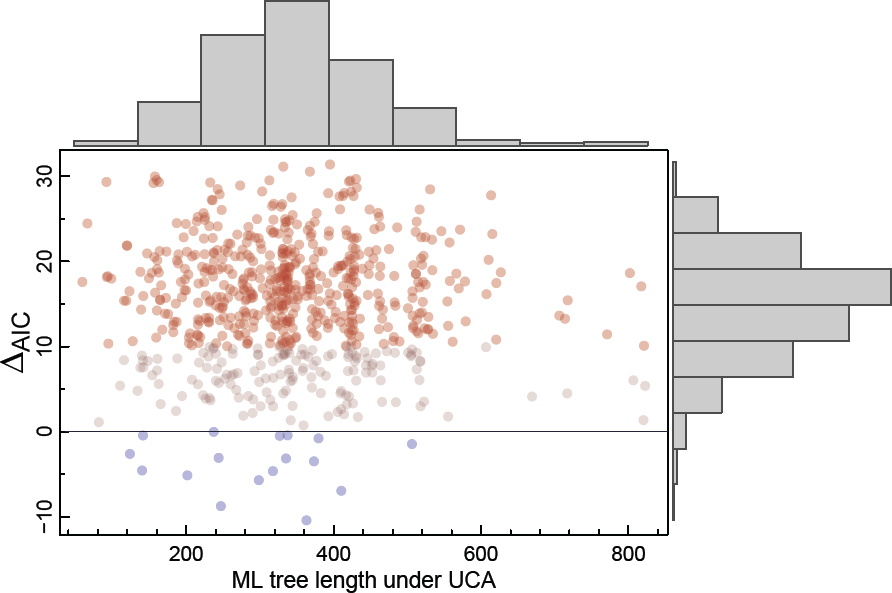
Δ*AIC* values for the simplified version of Koonin & Wolf’s simulations. Positive values for Δ*AIC* indicate UCA,and to ease interpretation the simulations favoring IO are displayed as blue dots, while those strongly favoring UCA (Δ*AIC* **>** 10) are red. Marginal histograms are also shown, and the gray dots represent simulations favoring UCA only slightly.

### 1.2 A note on pairwise versus phylogenetic comparisons

In his recent reply to K&W, Theobald gave arguments favoring Bayesian model selection over frequentist analysis [3]. But within his overview there seems to be some confusion about the advantages of his method against BLAST-based e-values: one thing are the virtues of Bayesian over frequentist analyses. Another, completely different issue, is the superiority of phylogenetic against pairwise sequence comparisons. His own solution to ”estimate an upper bound for the effect of alignment bias” was a random permutation of the sequences [3], which weakens his discourse against frequentist methodologies (as we will see this advice is incorrect since we cannot compare likelihoods between different data).

He compared a Bayesian phylogenetic model selection with a pairwise null hypothesis framework, praising the former over the latter [3]. But we would still prefer a *frequentist* phylogenetic model over a *pairwise* Bayesian one. That is, we might have a Bayesian model for pairwise homology detection [16] or a frequentist phylogenetic model selection, like for instance the one we present on Section 3. Likewise, we could devise a classic hypothesis testing where H0 and H1 are as described in the Appendix, using a branch length fixed at infinity against the alternative hypothesis with the length free to vary – we do not need to impose the same replacement matrix or other parameters across branches. In all cases the phylogenetic approach should be preferred over pairwise comparisons, because we expect that the effect of using the whole data at once should be more relevant than the statistical framework we choose.

## 2 Data sets conditioned on similarity

Theobald also suggested that his test works without corrections only for high confidence alignments [3], which we interpret as being those with low uncertainty and/or composed of similar sequences – this alignment quality requirement of the test was never ”formalized”. He mentioned ”eliminating any potential alignment bias”, where ’bias’ refers to ”artifactually induce[d] similarities between unrelated sequences” [3, page 14]. But to solve the UCA vs IO question we cannot restrict eligible data sets based on similarity, as by doing so we would be introducing an ascertainment bias towards alignments where UCA is more likely than for less similar ones. And notice that this is not to assume that similarity implies in homology, but it is a simple recognition that there is a correlation between them that cannot be neglected by excluding the sequences capable of refuting our hypothesis [7]. And we could even speculate that once the remove the ”alignment bias and uncertainty” what we are left with are columns that share a common ancestor *a fortiori*.

It could be finally argued that the test was designed instead only to distinguish UCA and IO from alignments that appear to favor UCA – that is, given similar-looking (or with elusively defined good quality) sequences, the test could detect independently originated sequences. However, Theobald himself didn’t bother about this constraint in his examples where the UCA test favored IO, as the experiment described in the last paragraph of page 221 and in the supplementary subsection 3.1 of [1]. In these experiments, alignment columns for a clade were randomly shuffled, resulting in very low pairwise similarity [4].

Theobald claimed that his test worked ”without assuming that sequence similarity indicates a genealogical relationship” [1], so we were interested in checking whether his test can indeed distinguish similar sequences with IO from similar sequences with an UCA. Indeed, it is hard to devise a simulation scenario where sequences generated under IO are very similar to each other, or are free from ”alignment bias”, and we have shown that all previous attempts failed at showing the correctness of the model [4]. We have argued that even summary statistics contain information about the likeliness of UCA, and therefore any common ancestry test must take this information into account [7]. Nonetheless it might be ultimately claimed that only bias-free alignments could invalidate the UCA test. Maybe a simulation where independent sequences should converge to a similar protein structure or to a limited set of structures might fit the demands, but we could not properly implement such a model at this point.

The closest approximation we could devise was to repeat the IO and UCA simulations as in [4], but now selecting the columns such that the average identity was above a given threshold. We must recall that this is not a proper simulation of highly similar IO sequences in general, since this toy example also suffers from a selection bias – and the frequency itself of column patterns defines a phylogeny [17, 18]. Specifically, we generated very long multi-sequence data sets under UCA or IO (as in other simulations [4, 7]), reordered their columns based on their conservation (from higher to lower average identity), and then selected exhaustively subsets of columns along this reordered data sets such that the average identity was above a threshold. We used segments of 1000 columns, which were each subjected to the UCA test twice: once before and once after aligning the segments with MUSCLE [19].

The results are shown in Figure 2, where we observe that the UCA hypothesis was always favored whenever the average sequence identity was higher than 0.44, even for sequences simulated under IO. Segments with similarities as low as 0.35 could also mislead the test in favor of UCA. And if we align the segments, then any sequences with more than 0.25 of average identity (before aligning) will be inferred as sharing a common ancestor, regardless of their actual relationship. Again, this is not an ideal simulation of highly similar IO sequences but still it suggests that by picking only the columns with high similarity we might falsely conclude for UCA. And importantly for our argument, it suggests that any reasonably conserved alignment would favor common ancestry no matter the actual origin of the sequences.

**Figure 2:**
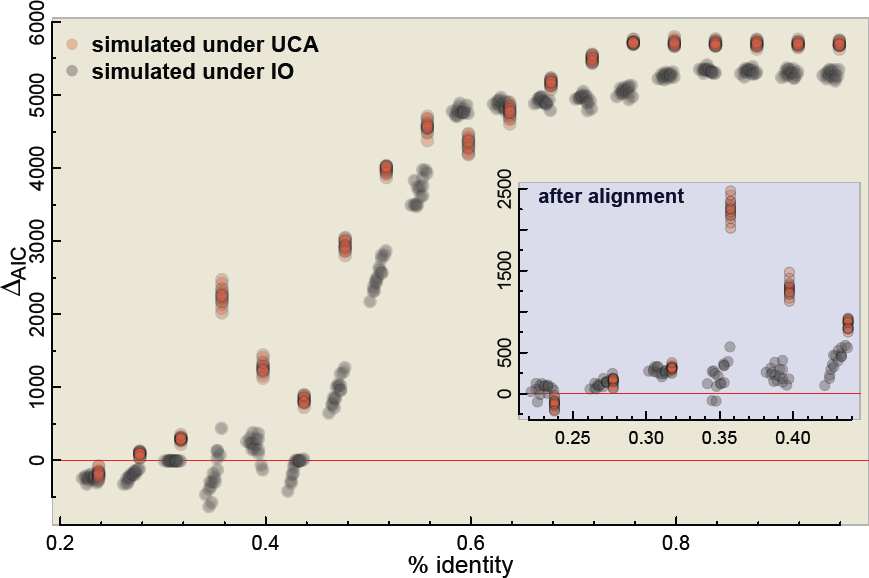
UCA test applied over large simulated data sets using a sliding window approach, where data sets’ columns were ordered from lower to higher average identity. Positive values for AIC suggest a UCA. The inset shows AIC after optimizing the alignment within the segments (where the average identity refer to the segment before the alignment step).

We could thus verify that the UCA test is oblivious to the source of the similarity: as long as the similarity is high enough it will favor UCA, while low similarities will have been previously camouflaged by an alignment optimization algorithm and even rejected altogether by BLAST or by the researcher, arbitrarily.

## 3 A random permutation test

An interesting alternative that does not rely on high quality alignments is to apply a permutation test where the sites for some sequences are shuffled and then the AICs are recalculated after realignment, telling us how much the original data departs from those with phylogenetic structure partially removed. It is inspired by [3, page 14], where it was suggested that the model selection test on one (or several?) shufflings would give an ”upper bound for the effect of alignment bias”. This randomization test has similarities to the permutation tail probability (PTP) tests [20, 21], but would invalidate the AIC and BF interpretations since the support value for UCA cannot be interpreted in isolation^3^. If we must compare the AICs between the original and randomized replicates – all favoring UCA, as we have shown –, then we are back to a frequentist analysis, where e.g. the AIC alone represents just a statistic that cannot be interpreted as probabilistic support for one of the hypotheses. Furthermore, we must emphasize that the AIC comparison only has a probabilistic interpretation when evaluated under the same data – the alignment, unless explicitly accounted for by the model (for phyml, prottest and others the data are the alignment columns). In other words, we cannot compare AICs between different alignments as suggested in [3], and even if we could, then this ”discount” should be an intrinsic part of any **formal** test. However, although the ”upper bound” argument is mistaken, it can lead to a valid permutation test.

This suggests that many other statistics may work in such a frequentist approach, that don’t need to rely on AIC or LnL values. We therefore developed such a randomization test where only simple statistics were considered, and applied it to *in silico* data sets. For each data set simulated under IO or UCA (same scenarios as in [4, Suppl. Mat.]) we calculate the statistics and then we create a distribution of these statistics under the hypothesis of independent origins (H0), to which the original value is compared (the p-value). Each H0 replicate is created by changing the columns order for one of the groups in the original data set, as was done in [1, section 3.1 of the suppl material] and described also in [4, Suppl. Mat. section S2.2]. Importantly, we always optimize the alignment for the original data set and each of the samples from H0 – so that we can estimate the ML tree, for instance. The statistics that we used are: 1) the sum of branch lengths of the ML tree for all sequences estimated under a LG model using phyML; and 2) the average pairwise identity within groups minus the average pairwise identity between groups suspected of having independent ancestry. In both cases we expect lower values for UCA than for IO, and our p-value is thus constructed by counting the number of null-distributed replicates presenting a value as low as the original data (where ”original data” is actually our data set simulated with INDELible).

In Figure 3 we show the results of 400 simulated data sets – 200 simulated under IO and 200 under UCA – where the null hypothesis was approximated by 100 shufflings (for each of the 400 data sets). We can see that not only the statistics are different between IO and UCA data sets, but that the p-values can clearly distinguish both cases (with the p-value uniform under the null, as expected). We could have used the AIC or BF scores from the original UCA test as the comparison statistics here, but it would give us similar results. And they would not give us any further insight, since their individual values would ”support” UCA even under IO [4] and even their differences or ratios would not represent statistical support anymore. Here we show again, as in [7], that the alignment properties are by themselves informative about UCA, and even without employing the whole AIC-based model selection analysis we can test for UCA. We should note that we do not endorse this test as the ultimate solution: as discussed by [21], the PTP test itself is flawed (but see [22]) and there might be caveats with our version as well. We suspect that our random permutation test would fail just like the original UCA test for bias free or ”very high confidence” alignments of IO sequences, but as we argue this is an unreasonable assumption to start with. The ”alignment bias” should not be used as a criterion for the adequacy of the UCA test, since it is an integral part of any **formal** common ancestry test.

**Figure 3:**
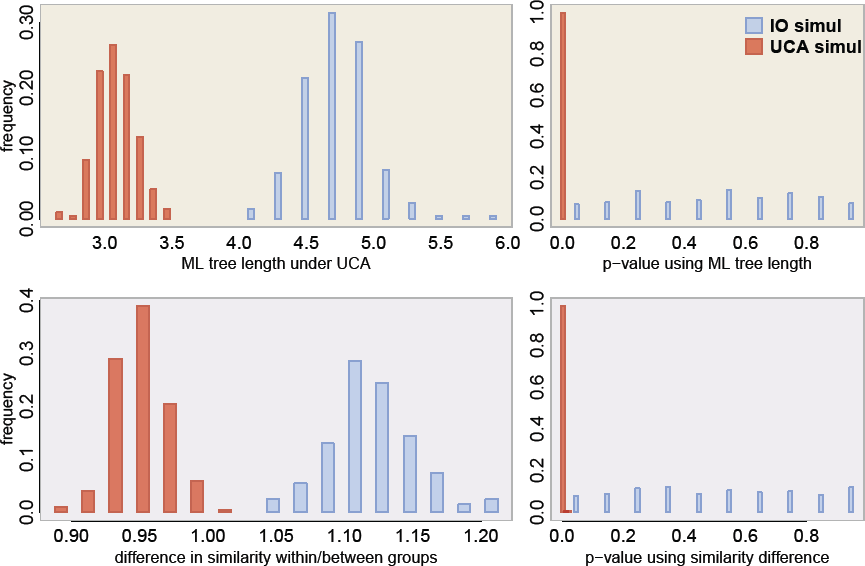
Frequentist nonparametric p-values where the null distribution was approximated by reshuffling columns of a subset of the sequences. On the left we have the distribution of the test statistics for the ”original” sequences simulated under IO (blue) or under UCA (red), while the right show their associated p-values. At the top the test statistic is the maximum likelihood tree length under the common ancestry hypothesis, and at the bottom the statistic is the difference in average similarity within each group and between one group and the other.

### 3.1 Aligning independently under each hypothesis

Model selection tests can help deciding between models for a given data set, but they cannot be compared across different data. Therefore we should not compare e.g. AIC values or log-likelihood ratios between different alignments, as under phyML and many other programs that do not consider explicitly the indel process. In contrast to e.g. [23], for these programs the data are the frequency of site patterns (i.e. the alignments). That is, alignment columns are considered independent and identically distributed observations from the evolutionary process. Therefore we stress that in order to apply the original UCA model selection test we must use the same alignment for both the IO and UCA hypotheses.

But what values would we observe if we could simply align the sequences independently, as has been suggested by Theobald [personal communication]? For this simulation we used the same simulation scenario as before [4] assuming an LG+IGF for each independently originated quartet – that we call B and E since they are based on bacterial and eukaryotic parameters, respectively. But now under the IO hypothesis we align the quartets separately – that is, in order to calculate the *AIC*(*B*) we align only the B sequences, and so forth. We can also try to account for the different alignment sizes by using the Bayesian Information Criterion (BIC) [24]:

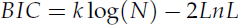

which is similar to the AIC but where we have the log of the number of data points (=column alignments in our case) instead of a fixed integer. The alignment size will be generally the same under each IO subset, and will correspond to the original sequence size of 6591 sites, while under UCA it will be around 10% larger, indicating the imputation of indels if we align both subsets together [4]. The Δ*AIC* values do not change whether we align the putative independent data sets together or separately, and trying to correct for the alignment size makes the tests perform even worse (Figure 4). Therefore, even if we align each subset independently from the others, we would still observe misleading, positive Δ*AIC*s. Again, the probabilistic interpretation of these information criteria is lost under this procedure.

**Figure 4:**
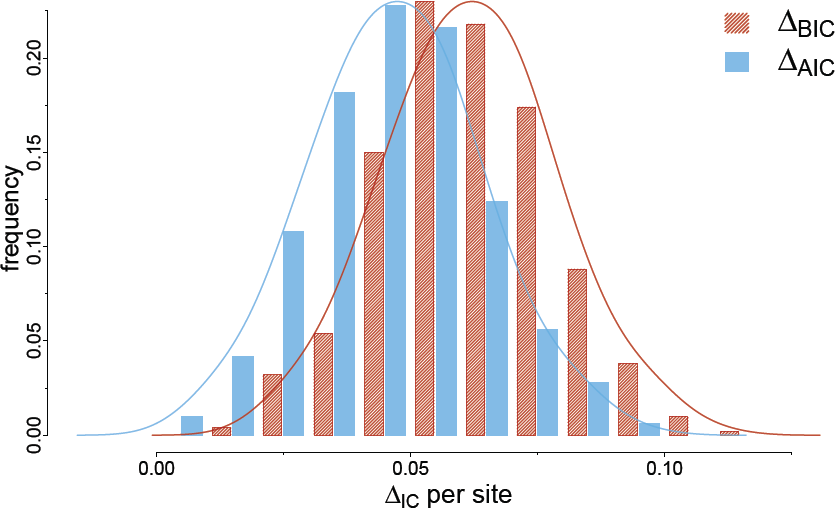
AIC and BIC per site if we align independently each subset (B and E sequences under IO, and B+E sequences under UCA). Each Δ*IC* (where IC=AIC or BIC) is calculated as Δ*IC* = *IC*(*B*) + *IC*(*E*) − *IC*(*B* + *E*) such that positive values favor UCA. The Δ*IC* values were further divided by the alignment length under UCA, to give scaled values comparable with other analyses.

## 4 Discussion

In Theobald’s response to K&W’s simulations, he showed that by extending his test to include the true model (the MAX-Poisson under a star tree with infinite branch lengths, called ”profile” model) it would be preferred over a single tree with a standard substitution model. This shows that the evaluated phylogenetic substitution models are consistent, but do not provide evidence about the appropriateness of the original UCA test. Even more, the actual model selection should be thought of as a blind test: we must not rely on some privileged knowledge about the true origin of the data set to reject hypotheses beforehand. Since we never know the true generating model of real data sets – which is specially true in phylogenetics – we must accept that all models we work with are misspecified [24].

On the other hand, if the inference for or against UCA depends on details of the phylogenetic model, then the test will only be useful when we know the true phylogenetic model. We do not expect a useful model to be very sensitive to model violations, specially when these violations can be assumed to affect both hypotheses. We expect the test to favor the correct hypothesis for any model close enough to what might be the true generating one.

For example if our conclusion for UCA or IO changes depending on whether allow or not rate heterogeneity, whether we include or not a given replacement matrix, or some other mild model misspecification, then it becomes hard to defend our conclusion, and we should not trust this model selection. Our expectation is that a model good enough will affect both hypotheses likewise.

We are not against extending the framework to include more models, which might help distinguishing an IO data set from an UCA one. After all, the test output will give the odds ratio given a set of assumptions – like for instance rate heterogeneity, common branch lengths along the alignment, a common topology for all sites, etc. And we can always improve on the assumptions. Furthermore if we can devise an evolutionary model whereby independent sequences can mislead BLAST searches and alignment procedures, certainly we would like to see it implemented it in such a model selection framework. But we should accredit it as a contribution to a better model selection test, specially if such model could have systematically misled the original one. Systematically misleading simulations are a valid criticism to a particular model selection scheme, that deserve credit.

We shouldn’t dismiss a model based solely on our subjective impressions about commonplace data sets, either: novel methodologies are created precisely to discover patterns that were hidden or unexplained so far. Therefore biological realism or representativeness may not be good judges of a model’s relevance. In exploratory analysis we employ several shortcuts like skipping similar models or disregarding those based on assumptions known to be very unlikely. But when the aim is to assign objectively probabilities to the hypotheses, then we should consider and embrace models capable of refuting them.

A more serious problem may be when model misspecification happens only under one of the hypothesis (due to software limitations, for instance). For instance, cases where amino acid replacement model heterogeneity between the independently evolved data sets can affect the test: while under UCA all branches are forced to follow the same replacement matrix, gamma parameter and equilibrium frequencies, under IO the independently evolved groups are allowed to have their own ones. We recognize that this is an implementation problem and not a theoretical one – programs usually make this homogeneity assumption to avoid overparameterization. Nonetheless, we should be careful whenever the test favors IO since it might be the case of a better parameterization – one set of parameters for each subtree. Whenever the test favors IO, we should always try to isolate the effect of the IO assumption against the confounding effect of amino acid replacement heterogeneity by one of two ways.

One is by extending the software to replace the fixed parameter by a variable one. That is, to allow the implemented model to have a variable replacement matrix along the tree, or a heterogeneous equilibrium frequency vector across branches, etc. so as the UCA tree can access the same parameter space as the IO trees. The other is to assume homogeneity under the IO hypothesis by using the same parameters over all independently evolved groups, such that any model misspecification can be ”marginalized”. If some apparent support for the IO hypothesis disappears once we force homogeneity, then we can suspect that the model misspecification was misleading the test.

We maintain that the UCA test as originally proposed [1] is heavily biased towards UCA, but a good counterargument would be to show a replicable simulation procedure that generates bias-free alignments where the test correctly detects IO. The problem lies in that there are no known mechanisms by which we can simulate independently evolved sequences that satisfy the quality requirements imposed in [3] – and any attempt might be met with a special pleading, as we have seen. It is worth noticing that another method has been recently proposed that can more directly test for ancestral convergence [12]. This method does not seem to suffer from the drawbacks of the UCA test, since it takes into account the alignment step.

Another powerful argument for the common ancestry of life is to show how distinct genes or different units of information support similar phylogenetic histories – and we thank Douglas Theobald for the herculean task of compiling the evidence for it in an accessible manner (<u>http://www.talkorigins.org/faqs/comdesc/</u>). But unfortunately the opportunity of showing this consilience of trees for the universally conserved proteins was missed: the UCA model selection framework suggested that several trees were much more likely than a single tree for all proteins [1], which *prima facie* goes against a universal phylogeny, in the absence of a quantification of the amount of disagreement. We are thus left only with a visual corroboration of the non-random clustering of taxa [1, Figure 2a], which do indeed provide evidence for the common ancestry of the analysed sequences.

1 equals one if and only if *χ* is true and equals zero otherwise

2 Although we must never expect real data sets to follow exactly an implemented model

3 We cannot compare likelihoods between different data

## APPENDIX – Independent origins as a special case of the common origin

Here we show in more detail our sketched proof that in the model selection proposed by [1] the IO hypothesis is a particular case of the model for UCA, if we assume a single evolutionary model *M* [4, Suppl Mat]. However, this conclusion remains valid if we relax the fixed model assumption: instead of a single evolutionary model we can think of variable models along the tree.

The following diagram represents how the UCA model (at the left) can lead to the IO model (at the right), where we see that after the “removal” of the internal branch the remaining neighboring branches have one less degree of freedom since the likelihood is the same whenever their sum is the same. In other words, for each independent origin three internal branches are fixed: one at ∞, representing the *de novo* appearance, and one at each of its sides becoming redundant by the pulley principle. The parameters *a* and *c* are constants between zero and one and a natural choice is *a* = *c* = 1, while A, B, C and D are subtrees (with one or more leaves).

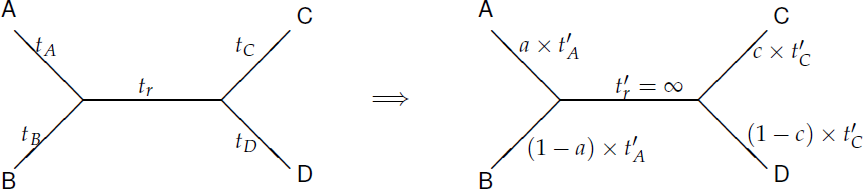

The justification for fixing the branch length at infinity comes from the fact that the Markov chains used in amino acid replacement models converge to their equilibrium distributions. That is, the probability *P*(*x | z*, *t*, *M*) of going from state *z* to state *x* in time *t* under evolutionary model *M* becomes independent of *z* when *t →* ∞, and approaches the equilibrium frequency *π*_*x*_ of *x* under model *M*:

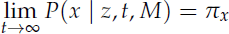

The likelihood *P*(*X | T*, **t**, *M*) of a phylogenetic tree *T* with branch length vector **t** arbitrarily rooted at *r* can be calculated for a given alignment column *X* as

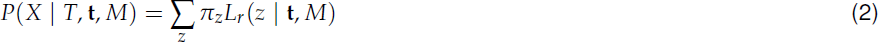

where *L*_*r*_(*z* | **t**, *M*) is the partial likelihood of node *r* for amino acid state *z*, and can be calculated recursively by

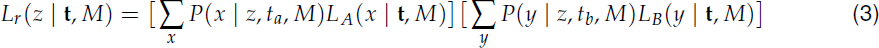

Assuming the subtree with branch lengths *t*_*a*_, *t*_*b*_ ∈ **t** connecting *r* to (internal or external) nodes *A* and *B* represented by

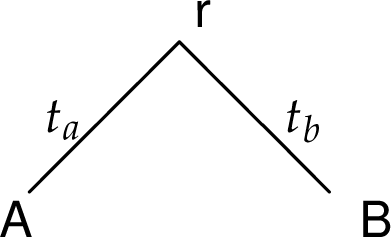

In the case when *t*_*a*_ and *t*_*b*_ go to infinity then as we saw *P*(*x | z*, *t*_*a*_, *M*) = *π*_*x*_ and *P*(*y | z*, *t*_*b*_, *M*) = *π*_*y*_, and therefore we have that equation 3 reduces to

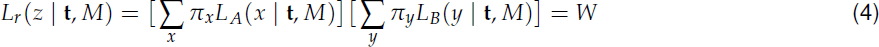

which is independent of *z*, and thus the site likelihood of equation 2 becomes

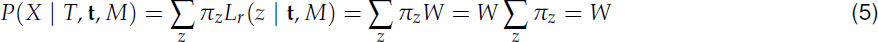

By comparing each of the two terms in equation 4 and equation 2 we can see that *W* is the product of the site likelihoods of two independent trees under the model described in [1], arbitrarily rooted at *A* and *B*. That is, if we represent the likelihood under the IO hypothesis as calculating independently the likelihoods of the subtrees rooted at *A* and *B* and multiplying them, then we have that each of these terms will be ∑*x π*_*x*_ *L*_*ρ*_(*x* | **t**, *M*) where *ρ* = (*A*, *B*), as in equation 2. The product of these terms is identical to the likelihood of a single tree with an infinite branch length connecting *A* and *B*, described by equation 5.

The extension for distinct amino acid replacement models over the tree is straightforward (replacing *M* by **M** = (*M*_1_, …, *M*_2*N−*2_) for *N* leaves), with the caveat that despite it can be handled by sequence simulation programs like INDELible [13], it is not implemented yet in popular phylogenetic reconstruction methods. Therefore the lack of correspondence of models *M* between the hypotheses is a limitation of the software employed, and not of the hypothesis test as devised.

## References

[1] Douglas L Theobald. A formal test of the theory of universal common ancestry. Nature, 465(7295):219–22, May 2010.

[2] Eugene V Koonin and Yuri I Wolf. The common ancestry of life. Biology direct, 5(1):64, January 2010.

[3] Douglas L Theobald. On universal common ancestry, sequence similarity, and phylogenetic structure: The sins of P-values and the virtues of Bayesian evidence. Biology direct, 6(1):60, November 2011.

[4] L. de Oliveira Martins and D. Posada. Testing for universal common ancestry. Systematic Biology, 63(5):838–842, Jun 2014.

[5] Takahiro Yonezawa and Masami Hasegawa. Was the universal common ancestry proved? Nature, 468(7326):E9; discussion E10, December 2010.

[6] Takahiro Yonezawa and Masami Hasegawa. Some problems in proving the existence of the universal common ancestor of life on Earth. The Scientific World Journal, 2012:479824, January 2012.

[7] Leonardo de Oliveira Martins and David Posada. Proving universal common ancestry with similar sequences. Trends in Evolutionary Biology, 4:e5, 2012.

[8] Nicolas Lartillot and Hervé Philippe. A Bayesian mixture model for across-site heterogeneities in the amino-acid replacement process. Molecular biology and evolution, 21(6):1095–109, June 2004.

[9] Le Si Quang, Olivier Gascuel, and Nicolas Lartillot. Empirical profile mixture models for phylogenetic reconstruction. Bioinformatics (Oxford, England), 24(20):2317–23, October 2008.

[10] Joseph Felsenstein. Evolutionary trees from DNA sequences: a maximum likelihood approach. Journal of molecular evolution, 17(6):368–76, January 1981.

[11] W J Bruno. Modeling residue usage in aligned protein sequences via maximum likelihood. Molecular biology and evolution, 13(10):1368–74, December 1996.

[12] W. Timothy J. White, Bojian Zhong, and David Penny. Beyond reasonable doubt: Evolution from dna sequences. PLOS ONE, 8(8):e69924, 2013.

[13] William Fletcher and Ziheng Yang. INDELible: a flexible simulator of biological sequence evolution. Molecular biology and evolution, 26(8):1879–88, August 2009.

[14] K.P. Burnham and D.R. Anderson. Model selection and multi-model inference: a practical information-theoretic approach. Springer, 2002.

[15] K. P. Burnham and David P. Anderson. Multimodel Inference: Understanding AIC and BIC in Model Selection. Sociological Methods & Research, 33(2):261–304, November 2004.

[16] Bobbie-Jo M Webb, Jun S Liu, and Charles E Lawrence. BALSA: Bayesian algorithm for local sequence alignment. Nucleic acids research, 30(5):1268–77, March 2002.

[17] MD Hendy and D Penny. A framework for the quantitative study of evolutionary trees. Systematic Biology, 38(4):297–309, 1989.

[18] David Bryant. Hadamard phylogenetic methods and the n-taxon process. Bulletin of mathematical biology, 71(2):339–51, February 2009.

[19] R.C. Edgar. MUSCLE: multiple sequence alignment with high accuracy and high throughput. Nucleic acids research, 32(5):1792, 2004.

[20] Daniel P. Faith and Peter S. Cranston. Could a Cladogram This Short Have Arisen By Chance Alone?: on Permutation Tests for Cladistic Structure. Cladistics, 7(1):1–28, March 1991.

[21] David L. Swofford, Jeffrey L. Thorne, Joseph Felsenstein, and Brian M. Wiegmann. The Topology-Dependent Permutation Test for Monophyly Does Not Test for Monophyly. Systematic Biology, 45(4):575, December 1996.

[22] Mark Wilkinson, Pedro R Peres-Neto, Peter G Foster, and Clive B Moncrieff. Type 1 error rates of the parsimony permutation tail probability test. Systematic biology, 51(3):524–527, 2002.

[23] J L Thorne, H Kishino, and J Felsenstein. Inching toward reality: an improved likelihood model of sequence evolution. Journal of molecular evolution, 34(1):3–16, January 1992.

[24] David Posada and Thomas R Buckley. Model selection and model averaging in phylogenetics: advantages of akaike information criterion and bayesian approaches over likelihood ratio tests. Systematic biology, 53(5):793–808, October 2004.

